# How the Central American Seaway and an ancient northern passage affected flatfish diversification

**DOI:** 10.1101/247304

**Authors:** Lisa Byrne, Franç Chapleau, Stéphane Aris-Brosou

**Affiliations:** Department of Biology, University of Ottawa, Ottawa, ON, CANADA; Department of Mathematics & Statistics, University of Ottawa, Ottawa, ON, CANADA

**Keywords:** Pleuronectiformes, Bayesian dating, trans-Arctic migration, vicariance, Central American Seaway, Isthmus of Panama

## Abstract

While the natural history of flatfish has been debated for decades, the mode of diversification of this biologically and economically important group has never been elucidated. To address this question, we assembled the largest molecular data set to date, covering > 300 species (out of *ca*. 800 extant), from 13 of the 14 known families over nine genes, and employed relaxed molecular clocks to uncover their patterns of diversification. As the fossil record of flatfish is contentious, we used sister species distributed on both sides of the American continent to calibrate clock models based on the closure of the Central American Seaway (CAS), and on their current species range. We show that flatfish diversified in two bouts, as species that are today distributed around the Equator diverged during the closure of CAS, while those with a northern range diverged after this, hereby suggesting the existence of a post-CAS closure dispersal for these northern species, most likely along a trans-Arctic northern route, a hypothesis fully compatible with paleogeographic reconstructions.

---

The Pleuronectiformes, or flatfishes, are a speciose group of ray-finned fish, containing 14 families and over 800 known species (Munroe 2015). Flatfish begin life in the pelagic zone, but undergo a larval metamorphosis in which one eye, either left or right, depending on the species, migrates to the other side of the cranium. The adult fish then adopts a benthic lifestyle. Flatfish have asymmetric, laterally-compressed bodies, and have lost their swim bladders during transformation. With eyes facing upwards, flatfish are also capable of protruding them. This singular morphology long puzzled taxonomists (Norman 1934), and the phylogeny of this group remains poorly resolved.

At the highest taxonomic level, flatfishes are generally considered to be monophyletic, based on both morphological (Chapleau 1993), and molecular evidence (Berendzen et al. 2002; Pardo et al. 2005; Azevedo et al. 2008; Betancur-R et al. 2013; Harrington et al. 2016). All these studies also support the monophyletic status of most families within the order, to the exception of the Paralichthyidae. As all molecular studies to date have essentially focused on the monophyletic status of the order, they were based on as many representative species of each order. As a result, intra-ordinal relationships, among genera and even families, are still debated. It can therefore be expected that taking advantage of both species- and gene-rich evidence, while incorporating paleontological and/or geological data in the framework of molecular clocks, should help clarify not only the phylogenetic status of this family (dos Reis et al. 2015), but also – and more critically here – their evolutionary dynamics.

However, very few flatfish fossils are known (Chanet 1997; Friedman 2012), and placed with confidence on the flatfish evolutionary tree (Parham et al. 2012; Campbell et al. 2014a; Harrington et al. 2016). As a result, calibrating a molecular clock becomes challenging. On the other hand, a dense species sampling may enable us to take advantage of a singular feature of flatfishes: some extant species are found both in the Pacific and Atlantic oceans. Furthermore, the existence of geminate species pairs of flatfishes, where sister taxa have one member in each ocean, suggests a speciation event pre-dating the formation of the Isthmus of Panama, which occurred approximately 12 to 3 million years ago [MYA] (Haug and Tiedemann 1998; O’Dea et al. 2016). Also possible is the role played by the opening of the Bering Strait around 5.5-5.4 MYA (Gladenkov et al. 2002), which would have permitted the trans-Arctic migration of ancestral populations from one ocean to the other, as documented in marine invertebrates (*e.g*., Durham and MacNeil 1967; Reid 1990) or vertebrates (*e.g*., Grant 1986), but to date, never in flatfish. Our driving hypothesis is then that the sole information relative to the formation of the Isthmus of Panama can be used to calibrate molecular clocks, and allows us to unravel the timing of flatfish evolution, as how rapidly they diversified remains an unsolved question. We show here that the diversification of flatfish in the seas surrounding the Americas followed a complex pattern, related to not only the closure of the Isthmus of Panama, but also to a warming event that opened up a trans-Arctic northern route.

## Results

To test how the evolutionary dynamics of flatfish were affected by a major geological event, the closure of the Central American Seaway (CAS), we reconstructed dated Bayesian phylogenetic trees from a large data matrix under four relaxed molecular clock models, each one of them being based on a different calibration scheme (Fig. 1). Under the first model, no calibration priors were placed on internal nodes. This initial tree, with the rogue sequences removed (data on GitHub: see Methods), was used to identify pairs of sister taxa that are split between the two oceans, with one species in the Atlantic and the other in the Pacific. This led us to single out twelve pairs of such species. These sister species happened not to be evenly distributed on the estimated phylogeny (Fig. 2, top), but they are the only geminate species included in GenBank (as of August 2016). On each pair, we placed calibration date priors corresponding to the closure of the CAS (the ‘ALL’ model). An examination of the posterior distributions of their divergence times suggests that some species have a very narrow speciation window, where all the mass of the posterior distribution is between 5-3 MYA, while others have a wider distribution (Fig. 3A). Closer inspection of these distributions further reveals that most of the species with narrow posterior distributions have a northern range (Fig. 2, and 3A, in blue), while those with the wider posterior distributions have a “southern” distribution, closer to the Isthmus of Panama (Fig. 2, and 3A, in red). To further assess this observation, we first went back to the original clock model, removed all priors initially placed on sister taxa (the ‘NONE’ model), and were able to validate that even in this case: northern and southern species showed, to one exception each (*Hippoglossus hippoglossus* and *H. stenolepis* in the north, and *Poecilopsetta natalensis* and *P. hawaiiensis* in the south; Fig. 2), shifted posterior distributions (Fig. 3B). The former pair was actually not estimated as being sister species in any of the four clock models (Fig. 1), while *P. natalensis* and *P. hawaiiensis*, although inhabiting the Western Indian and Eastern Pacific oceans, occupy ranges that do not extend to the coasts of the Americas as with the other identified sister taxa. Models with priors placed only on northern (the ‘NORTH’ model: Fig. 3C) or southern (the ‘SOUTH’ model: Fig. 3D) species also showed a similar temporal shift. This shift suggested that southern species diverged early, before the complete closing of the CAS, while northern species diverged later, at or possibly after the isthmus was completed. Averaging these posterior distributions for the northern and southern species, to the exception of the two outliers noted above, showed these results more clearly (Fig. 3E-H). To assess the robustness of this result to the inclusion of the *P. natalensis* and *P. hawaiiensis* sister species, we conducted an additional set of analyses removing all prior information from this pair. The ALL and SOUTH models, which are the only ones where this prior occurs, show that the general pattern that we found above holds: southern species diverged early, before the complete closing of the CAS, and northern species diverged later, at or possibly after the isthmus was completed (Fig. S1).

**FIG. 1.**
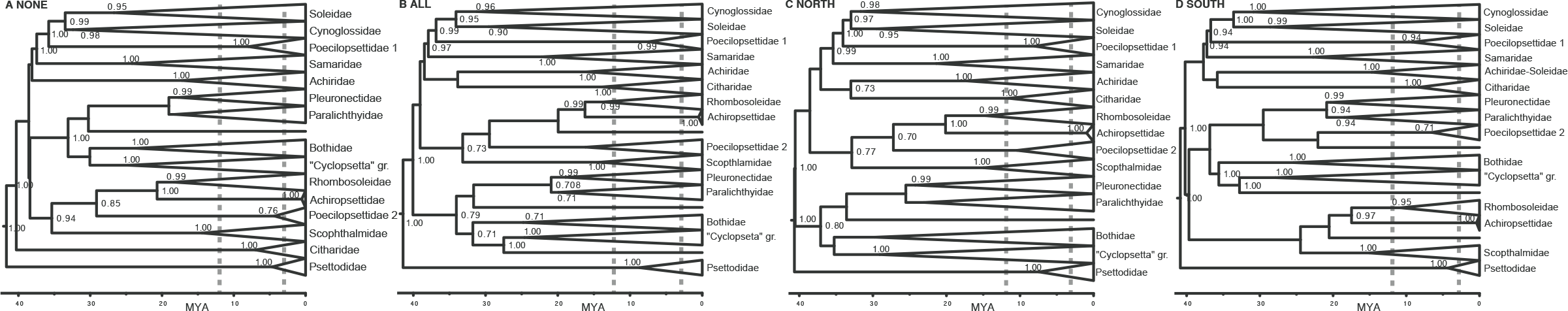
Phylogenetic trees reconstructed based on relaxed clock models. Four models were employed, representing different specifications of prior distributions set on sister taxa. (**A)** No priors were set on sister taxa. (**B)** Priors were set on all pairs of taxa. (**C)** Priors were set only on sister taxa with a current northern range. (**D)** Priors were set only on sister taxa with a current southern range. Horizontal scale is in million years ago (MYA). The closure of the CAS (12-3 MYA) is shown between vertical gray broken lines.

**FIG. 2.**
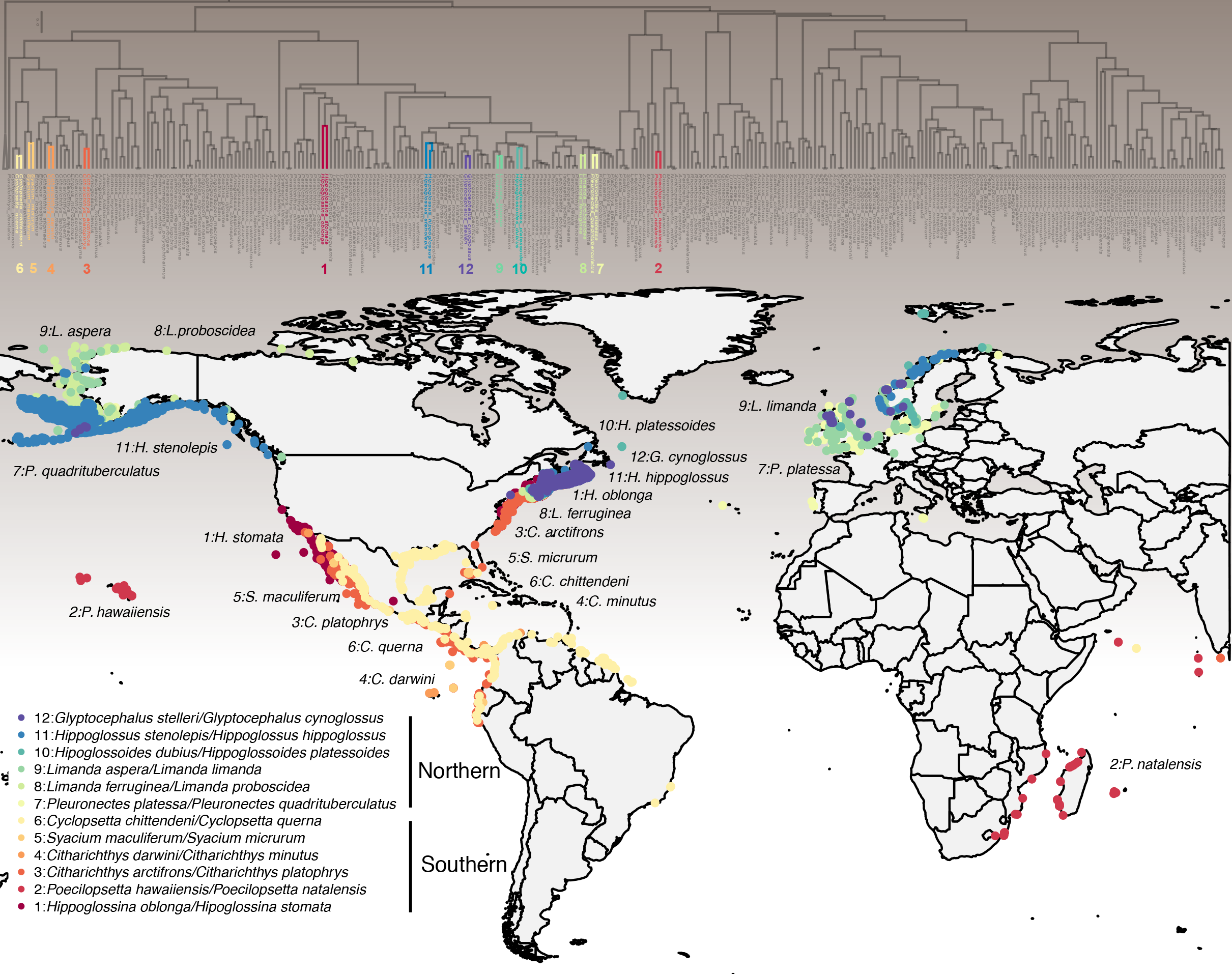
Distribution of the geminate species used in this study. Geographic distribution of the twelve pairs of sister species of flatfish used to calibrate the relaxed molecular clock models. Original data come from GBIF (www.gbif.org; accessed Nov 9, 2017). The top panel shows the phylogenetic distribution of these species based on the clock model including prior ages on all twelve pairs of species; scale bar: 5 MYA.

**FIG. 3.**
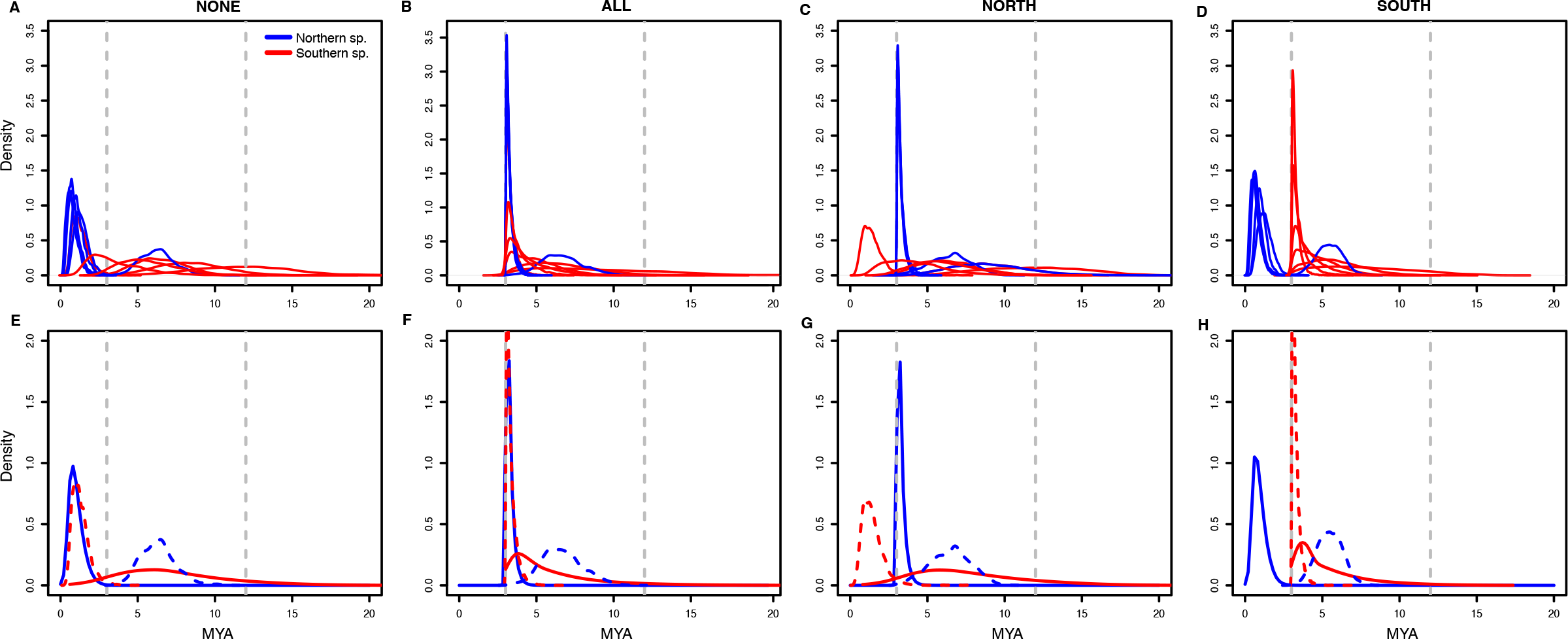
Posterior densities of divergence times for sister taxa used as calibration points in the relaxed molecular clock analyses, under four different calibration schemes. NONE: no priors were placed on sister taxa; ALL: priors were placed on all pairs of sister taxa; NORTH: priors were placed only on pairs of sister taxa with a northern distribution; SOUTH: priors were placed only on pairs of sister taxa with a southern distribution. Top row (**A**-**D**) shows all 17 distributions, while bottom row (**E**-**H**) shows range-averaged distributions (solid) to the exception of outlier pairs (dashed lines). Densities are color-coded for species with northern (blue) and southern (red) range. Dashed vertical gray lines indicate the closure of the CAS (21-3 MYA).

All these results were obtained assuming that Psettodidae is an appropriate outgroup for these analyses. However, the position of Psettodidae in flatfishes is still debated (Betancur-R. and Orti 2014; Campbell et al. 2014a; Campbell et al. 2014b; Harrington et al. 2016), so that their inclusion might affect our results. To assess this possibility, we reran all four models without Psettodidae, letting the molecular clock root the tree (Aris-Brosou and Rodrigue 2012). Figure S2 shows the exact same results as above (Fig. 3), so that our dating results are also robust to the inclusion, or not, of Psettodidae. Our dating results are also robust to the inclusion of internal calibration priors as used in Harrington et al. (2016), and based on the *Scophthalmus rhombus* / *Bothus pantherinus* and the *Crossorhombus kobensis* / *Bothus pantherinus* divergences, dated at 29.62 (Vandenberghe et al. 2012) and 11.06 MYA (Carnevale et al. 2006), respectively (Fig. S3).

In an attempt to tease out these models (including Psettodidae and without the two internal calibration priors) and their predictions about the exact timing of divergence between northern and southern species, we assessed model fit by means of the modified Akaike’s Information Criterion (AICM; Baele et al. 2013), which accounts for uncertainty in the MCMC sampling (Raftery et al. 2007). Even if model ranking based on this measure is known to be unstable (Baele et al. 2013), it is clear that the models with priors only on the northern or on the southern species perform significantly (> 200 AIC units) worse than the two other models, which may be difficult to tease apart (Table 1). The predictions of the best models suggest that *H. hippoglossus* and *H. stenolepis*, both northern species, consistently diverged before the complete closure of the seaway, in tandem with the average southern species, and that the average northern species diverged in tandem with *P. natalensis* and *P. hawaiiensis* (Fig. 3E-F).

**Table 1.** AICM values for the phylogenetic analyses using four different calibration schemes. Models are ranked from the best AICM model (top) to the worst model. SE: standard error.

| Calibration Model | AICM value | SE |
| --- | --- | --- |
| ALL | 374,082.199 | +/-1.579 |
| NONE | 374,110.488 | +/-3.789 |
| SOUTH | 374,367.749 | +/-0.696 |
| NORTH | 374,746.628 | +/-3.789 |

One intriguing result from these analyses is that the northern sister species seem to come from a single clade (Fig. 2), which would suggest that it is possible to identify the direction of the trans-Arctic migration. For this, we retrieved the current species range of all flatfish analyzed here (Fig. S4), and indicated their average latitude on the phylogenetic trees (Fig. S5). This northern clade, highly supported (posterior probability > 0.9; see asterisk in Fig. S5) even in the analyses including the two additional internal priors (Fig. S6), is mostly comprised of species currently found in the Pacific (*e.g*., *Reinhardtius evermanni*), suggesting that the northern sister species that we identified originated from the Pacific, and that their trans-Arctic migration was from the Pacific into the Atlantic in a northern direction.

## Discussion

Our results show that flatfish underwent a first speciation at the time when the CAS closed, which led to the species that, today, have an equatorial range (Fig. 2), as the formation of the Isthmus of Panama resulted in a barrier to gene flow leading to their speciation. Our results also show that species that today have a northern range (Fig. 2) either emerged at the closure of the seaway, or after its closure. These timing estimates imply that this second bout of speciation was not caused by gene flow impeded by the closure of the seaway, but demand an interpretation involving trans-Arctic gene flow, through a northern route, before being interrupted, most likely this time by a climatic event. Critically, these conclusions are robust to the inclusion of outgroup species (Psettodidae), of outliers species (*P. natalensis* and *P. hawaiiensis*), and are consistently found across the four clock models implemented here, including the one (NONE) relying on no other fossil information than the 47.8-42.1 MYA ancestor to this order (Chanet 1997; Friedman 2012) – hereby indicating that our data are highly informative with respect to the timing of the divergence of these sister species (and rejecting our initial hypothesis) following their trans-CAS and trans-Arctic migrations.

Trans-Arctic migrations from the Pacific to the Atlantic are not new, and have been described in both invertebrates (*e.g*., Durham and MacNeil 1967) and vertebrates (*e.g*., Grant 1986), but never using a relaxed molecular clock relying on as little information as a single fossil and a large multigene data set covering an entire order (Pleuronectiformes). While seminal work was usually based on paleontological (Durham and MacNeil 1967), or on morphological evidence (Reid 1990), subsequent work started using allozymes and other proteins (Grant 1986; Zaslavskaya et al. 1992), followed by genetic evidence, mostly at the population level using mitochondrial DNA (Rawson and Harper 2009). However, most of the recent work that relies on molecular clocks either makes strong assumptions on substitution rates (Rawson and Harper 2009; Liu et al. 2011), uses divergences that are vicariance-motivated but not supported by any fossil information (McCusker and Bentzen 2010), or posit in which direction the trans-Arctic migration took place (Dodson et al. 2007) to infer trans-Arctic migrations. Our work is the first to use sister species to demonstrate the existence and elucidate the timing of both the trans-CAS and the trans-Arctic events, while also inferring in which direction the latter took place, without making too many assumptions.

Strikingly, the geological evidence is directly in line with our date estimates. The fossil record suggests that the first aquatic connection between the Pacific and Arctic (and Atlantic) oceans through the Bering Strait occurred approximately 5.5-5.4 MYA (Gladenkov et al. 2002) due to a rise in sea levels, linked to tectonic activity (Marincovich and Gladenkov 2001). This would have permitted the trans-Arctic migration of populations ancestral to today’s northern species from one ocean to the other through this ancient “northern passage.” This global warming event, between the late Miocene to early Pleiocene, was then followed by a significant period of cooling during the Pleiocene into the Pleistocene (Zachos et al. 2001), leading to periods of repeated glaciations and a subsequent ice age. These cold events would have resulted in the closure of this ancient “northern passage,” hereby stopping the trans-Arctic gene flow between the two oceans, and leading to the more recent speciation of the northern taxa.

For approximately a million years after the first opening of the Bering Strait, water flowed through the strait in a southern direction, from the Arctic to the Pacific ocean, until the formation of the Isthmus of Panama occurred close to the equator (Berta 2012). With the formation of the Isthmus, and hence the closing of the CAS, the ocean currents reversed due to a change in global ocean circulation (Haug and Tiedemann 1998; De Schepper et al. 2015), and have since flowed in a northern direction through the Bering Strait, from the Pacific to the Arctic (Marincovich 2000). This again is absolutely consistent with our results (Fig. S4), with dispersal or migration from the Pacific into the Atlantic in a northern direction through this strait, which is known as the trans-Arctic interchange (Vermeij 1991). Fossil data also show that the Bering land bridge has been exposed and submerged on multiple occasions since the Pleistocene (Gladenkov and Gladenkov 2004). These openings and closings of the Bering Strait could have provided a mechanism for divergence and the evolution of sister taxa (Taylor and Dodson 1994; Väinölä 2003).

Our results also have implication at the family level of flatfish. Further significant global cooling during the Pleistocene resulted in major glaciation events (Zachos et al. 2008) that could be responsible for creating barriers that isolated populations. All of the remaining sister taxa in our analysis, who have divergence estimates of less than 2 MYA in our study belong to the family Pleuronectidae (*sensu* Cooper and Chapleau 1998). The Pleuronectidae are the predominant flatfish family found in cold and temperate seas of the northern hemisphere (Norman 1934; Cooper and Chapleau 1998). There are far more Pleuronectidae species in the Pacific Ocean, most of them endemic to the north Pacific Ocean off of North America and Asia in the region extending from the Bering Strait to the gulf of California (Norman 1934). None of the arctic species are restricted solely to arctic waters (Munroe 2015). Munroe also noted that Cooper (in an unpublished manuscript) identified areas of endemism among the current distribution of the Pleuronectidae. It has been shown that during the trans-Arctic interchange, there was a far higher number of species (up to eight times higher) migrating to the Atlantic than to the Pacific (Vermeij 1991). Fossil and phylogenetic evidence suggest the Pacific Ocean as the origin for diversification of the Pleuronectidae (Munroe 2015), and our phylogenetic results are highly congruent with this hypothesis.

It is possible that the outliers, the Atlantic and Pacific halibut (*H. hippoglossus* and *H. stenolepis*, respectively), diverged during one of the first openings of the Bering Strait. The estimated dates from the molecular clock analysis are approximately 5.5 MYA, in accordance with the hypothesized dates for the first aquatic connection (Marincovich and Gladenkov 2001). The remaining sister taxa have a younger age estimate of ~1-2 MYA, corresponding with global cooling during the Pleistocene and repeated glaciations (Zachos et al. 2001), which likely formed more barriers to genetic flow. In the second pair of outliers, *P. hawaiiensis* inhabits waters of the Eastern Central Pacific to the Hawaiian Islands, while *P. natalensis* inhabits coastal waters of Eastern Africa (Fig. 2). Due to the far reaching range of *P. natalensis*, and the relatively younger age estimates for divergence (1-2 MYA), these results beg for future research into other vicariant hypotheses, or dispersal routes, as these two members of *Poecilopsetta* diverged long after the Isthmus had closed. One candidate would be the Indo-Pacific barrier, which formed an almost continuous land bridge between Australia and Asia, and could have hence limited dispersal between the Pacific and the Indian oceans, hereby promoting speciation (Gaither et al. 2011). This barrier formed during the Pleistocene glacial cycles (2.58-0.01 MYA; Voris 2000), which is, again, fully consistent with our date estimates (Fig. 3, S1).

Based on the most extensive multigene sequence alignment available to date across all flatfish species, we have showed here that the evolutionary dynamics of sister species that are distributed across the two oceans strongly supported the existence of two bouts of speciation: one triggered by the closure of the CAS 12-3 MYA, and a second one due to the closure of an ancient northern passage 5-0.01 MYA. Our work adds to a growing body of evidence pointing not only to the role of geological processes in shaping biotic turnovers (Bacon et al. 2015), but also on the consequences of climate change that can either trigger speciation events (Reid 1990), or promote faunal interchanges whose ecological and economic impacts are difficult to predict (Wisz et al. 2015).

## Materials and Methods

### Data retrieval and alignment

Prior to retrieving sequence data, GenBank was surveyed to identify all the genes belonging to species of Pleuronectiform families (as of August 2016), based on GenBank’s taxonomy browser. DNA sequences for a total of 332 flatfish species (out of over 800 species in the order) were identified and downloaded for five nuclear genes (KIAA1239, MYH6, RIPK4, RAG1, SH3PX3), and four mitochondrial genes (12S, 16S, COX1 and CYTB). These represented all the taxa having at least one of these six gene sequences in GenBank; see Table S1 for the corresponding accession numbers. Diversity was richly sampled as species from 13 of the 14 families in the order Pleuronectiformes were included in our catchment. In line with current consensus in flatfish systematics, the family Psettodidae was chosen as the outgroup to all other taxa (Chapleau 1993).

These sequences were aligned using MUSCLE ver. 3.8.31 (Edgar 2004) on a gene-by-gene basis. Each alignment was visually inspected with AliView ver. 1.18 (Larsson 2014), and was manually edited where necessary. In particular, large indels (> 10 bp) were removed prior to all phylogenetic analyses. The 5’ and 3’ ends of sequences were also trimmed. The aligned sequences were then concatenated into a single alignment using a custom R script.

### Data pre-processing

To gauge the phylogenetic content of our data set, we performed a first series of molecular clock analyses on the concatenated data matrix, with all the nine gene sequences obtained above (12S, 16S, COI, Cyt-b, KIAA1239, MYH6, RIPK4, SH3PX3 and RAG1), and all the 332 taxa representing all families of flatfish. To account for rate heterogeneity along this alignment, partitions were first identified with PartitionFinder 2 (Lanfear 2012), selecting the best-fit model of nucleotide substitution based on Akaike’s Information Criterion (AIC). This optimal partition scheme was then employed in a first Bayesian phylogenetic analysis, conducted with BEAST ver. 1.8.3 (Drummond and Rambaut 2007). This program implements a Markov chain Monte Carlo (MCMC) sampler, which co-estimates both tree topology and divergence times. As determined by PartionFinder, a GTR+I+Γ model was applied to each data partition. These models of evolution across partitions were assumed to be independent (*unlinked*, in BEAST parlance), while both clock model and tree model partitions were shared (*linked*) by all partitions. An uncorrelated relaxed clock was employed with a lognormal distribution prior on rates, and a Yule speciation prior (Drummond et al. 2006). Due to the paucity of reliably placed fossils on the flatfish tree, a unique calibration point was placed on the most recent common ancestor (MRCA) of the ingroup, as a mean-one exponential prior, with an offset of 40 million years reflecting the age of 47.8-42.1 MYA (Chanet 1997; Friedman 2012). To stabilize the estimate, a lognormal ln(0, 1.5) prior with a 40 MYA offset was also placed on the root of the tree (root height). Two separate MCMC samplers were run, each for 10,000,000 generations. Trees were sampled every 5,000 generations, and convergence was checked visually using Tracer ver. 1.6 (available at: http://tree.bio.ed.ac.uk/software/tracer/). Tree log files from each run were combined in LogCombiner, after conservatively removing 10% of each run as burn-in. The resulting maximum a posteriori (MAP) tree was generated with TreeAnnotator (Drummond and Rambaut 2007).

As the topology of this resulting MAP tree was unconventional, we suspected the presence of rogue taxa, *i.e*. species evolving either much faster or much slower than the majority, which can contribute negatively to consensus tree support (Aberer et al. 2013). Rogue taxa were identified using the RogueNaRok (Aberer et al. 2013) webserver (http://rnr.h-its.org). The consensus trees from the preliminary analysis using 10,000 iterations were used as the tree set for rogue taxon analysis. A total of 22 rogues were identified and pruned from the analysis, leaving 310 species.

### Molecular Dating

To assess the impact of the closure of the CAS on flatfish evolutionary dynamics, a second set of partitioned relaxed molecular clock analyses was performed (without the rogue sequences). The timing of the closure of the CAS is estimated to have occurred between 12 and 3 MYA (Duque-Caro 1990; Coates et al. 1992; Haug and Tiedemann 1998; O’Dea et al. 2016), and we used this time window as a prior to inform the relaxed molecular clock-based phylogenetic reconstructions. The analyses were performed on the same concatenated data set, with the same single fossil calibration, but we also placed a lognormal prior ln(3,1.5), that has most of its mass on the 12-3 MYA window, on the MRCA of each pair of sister taxa (the ‘ALL’ model). From the initial BEAST analyses, twelve pairs of taxa were selected based on two criteria: (i) being sister species on that initial tree, (ii) with one species distributed in the Pacific and one in the Atlantic ocean (Fig. 2). Again, two independent MCMC samplers were run each for 100 million iterations, with samples taken every 5000 step.

Because these pairs of sister species show a contrasted geographic distribution, having either a southern (equatorial) or a northern range (Fig. 2), two additional sets of analyses were performed. In a first set, calibration priors (ln(3,1.5), as *per* above) were placed only sister taxa that had a geographic range in the southern hemisphere (the ‘SOUTH’ model), while in a second set, identical calibration priors were placed only on sister species with a northern range (the ‘NORTH’ model). Finally, a set of analyses was performed using no sister taxa calibrations at all (the ‘NONE’ model). For each analysis, results from the two MCMC runs were combined using LogCombiner after removing an even more conservative burn-in of 50%. The final MAP tree was generated with TreeAnnotator.

In an attempt to rank these different models (priors on all sister taxa; only on southern taxa; only on southern taxa; no “CAS” priors), the AICM was computed for each model (Baele et al. 2013). These computations were performed in Tracer for each of the four different calibration models, based on 100 replicates.

### Species distribution data

In order to map the distribution of each species of flatfish, we resorted to the Global Biodiversity Information Facility (GBIF: https://www.gbif.org/) database, accessed with the R library rgbif (Chamberlain 2017). Up to 1,000 records were retrieved for each species, and plotted on a geographic map based on GBIF observational records. Where needed, species locations were summarized by mean latitude and longitude (Fig. S4). Approximate attribution to a particular ocean was based on species-specific mean longitudes (Atlantic: between 20˚ and 120˚; Indian: 120˚ and −100˚; Pacific otherwise).

### Availability of Computer Code and Data

Accession numbers, sequence alignments, BEAST models (as xml files), estimated phylogenetic trees (maximum *a posteriori* trees from BEAST), and the R scripts used to plot the figures in this study are available from https://github.com/sarisbro.

## Acknowledgments

This work was supported by the University of Ottawa (LB), and the Natural Sciences Research Council of Canada (FC, SAB). We are grateful to Jonathan Dench for discussions and comments, to two anonymous reviewers for very constructive comments, and to Compute Canada and Ontario’s Center for Advanced Computing for providing us access to their resource. This work was completed while SAB was hosted by Yutaka Watanuki, at the University of Hokkaido in Hakodate, thanks to an Invitational Fellowship from the Japanese Society for the Promotion of Science.

