## Supplementary Materials for "How the Central American Seaway and an ancient northern passage affected flatfish diversification" (from another version of this preprint: version 2, 2018-03-26)

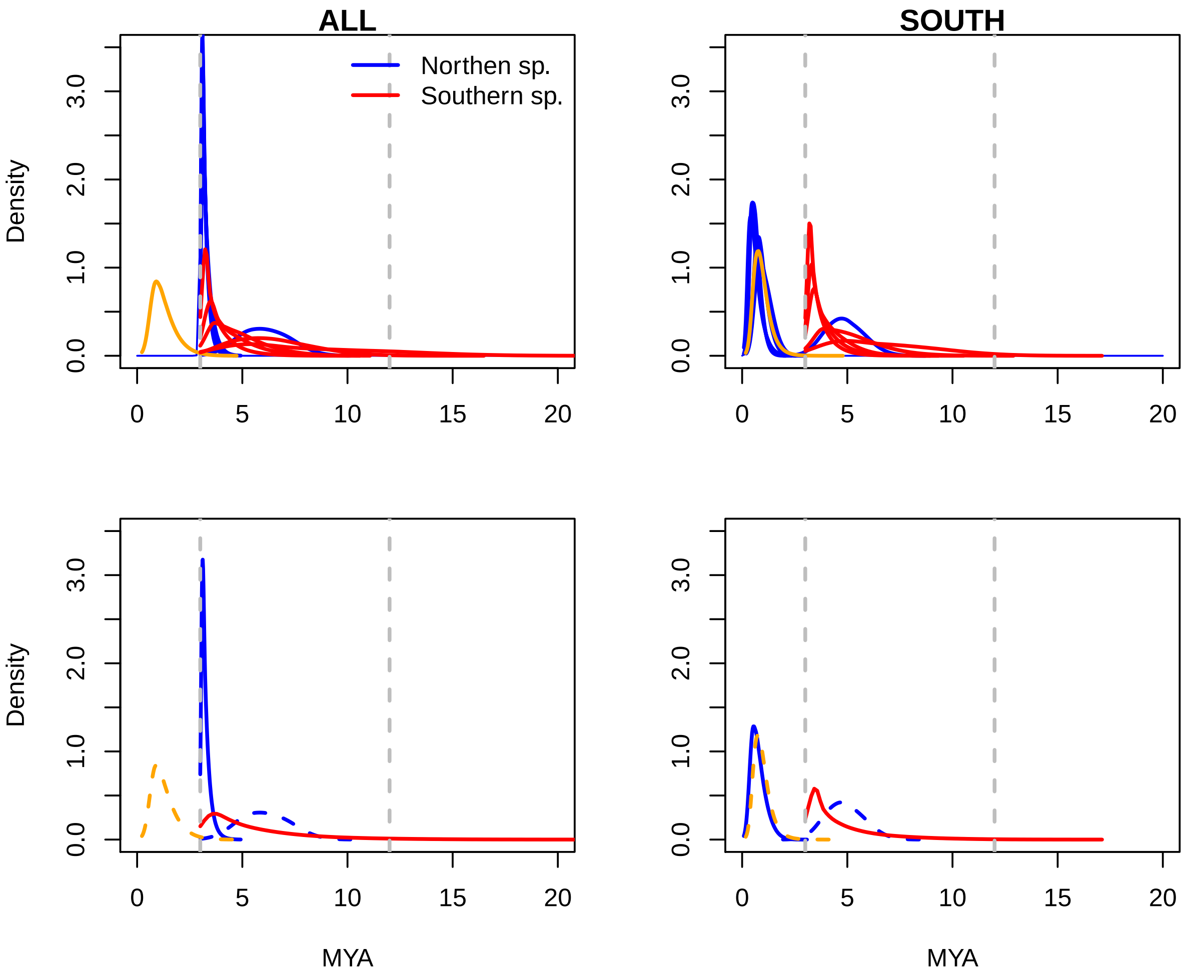


**Figure S1.** Posterior densities of divergence times for sister taxa used as calibration points in the relaxed molecular clock analyses excluding priors on *Poecilopsetta natalensis* and *P. hawaiiensis*. ALL: priors were placed on all the pairs of sister taxa; SOUTH: priors were placed on the pairs of sister taxa with a southern distribution. Top row shows all 17 distributions, while bottom row shows range-averaged distributions (solid) to the exception of outlier pairs (dashed lines; *Poecilopsetta natalensis* and *P. hawaiiensis* shown in orange). Densities are color-coded for species with northern (blue) and southern (red) range. Dashed vertical gray lines indicate the closure of the CAS (21-3 MYA).

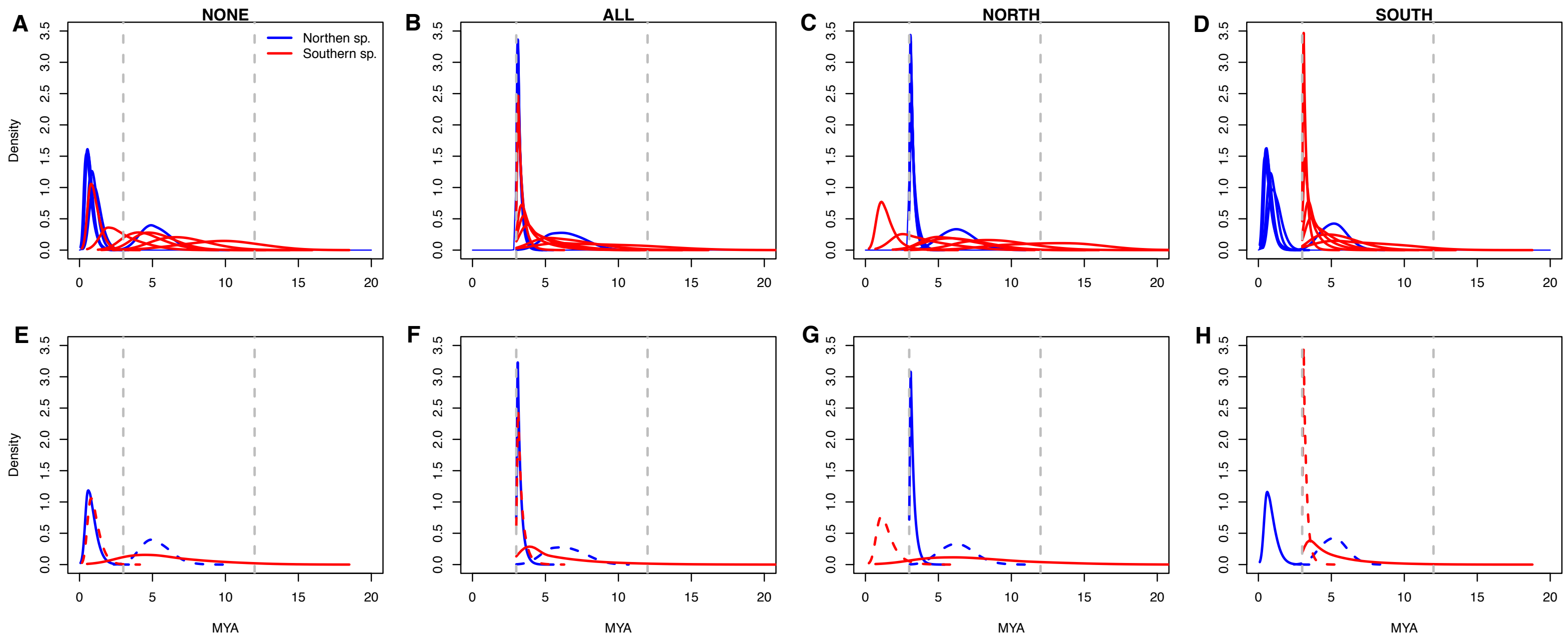


**Figure S2.** Posterior densities of divergence times for sister taxa used as calibration points in the relaxed molecular clock analyses excluding Psettodidae, under four different calibration schemes. NONE: no priors were placed on sister taxa; ALL: priors were placed on all pairs of sister taxa; NORTH: priors were placed on only pairs of sister taxa with a northern distribution; SOUTH: priors were placed on only pairs of sister taxa with a southern distribution. Top row (**A**-**D**) shows all 17 distributions, while bottom row (**E**-**H**) shows range-averaged distributions (solid) to the exception of outlier pairs (dashed lines). Densities are color-coded for species with northern (blue) and southern (red) range. Dashed vertical gray lines indicate the closure of the CAS (21-3 MYA).

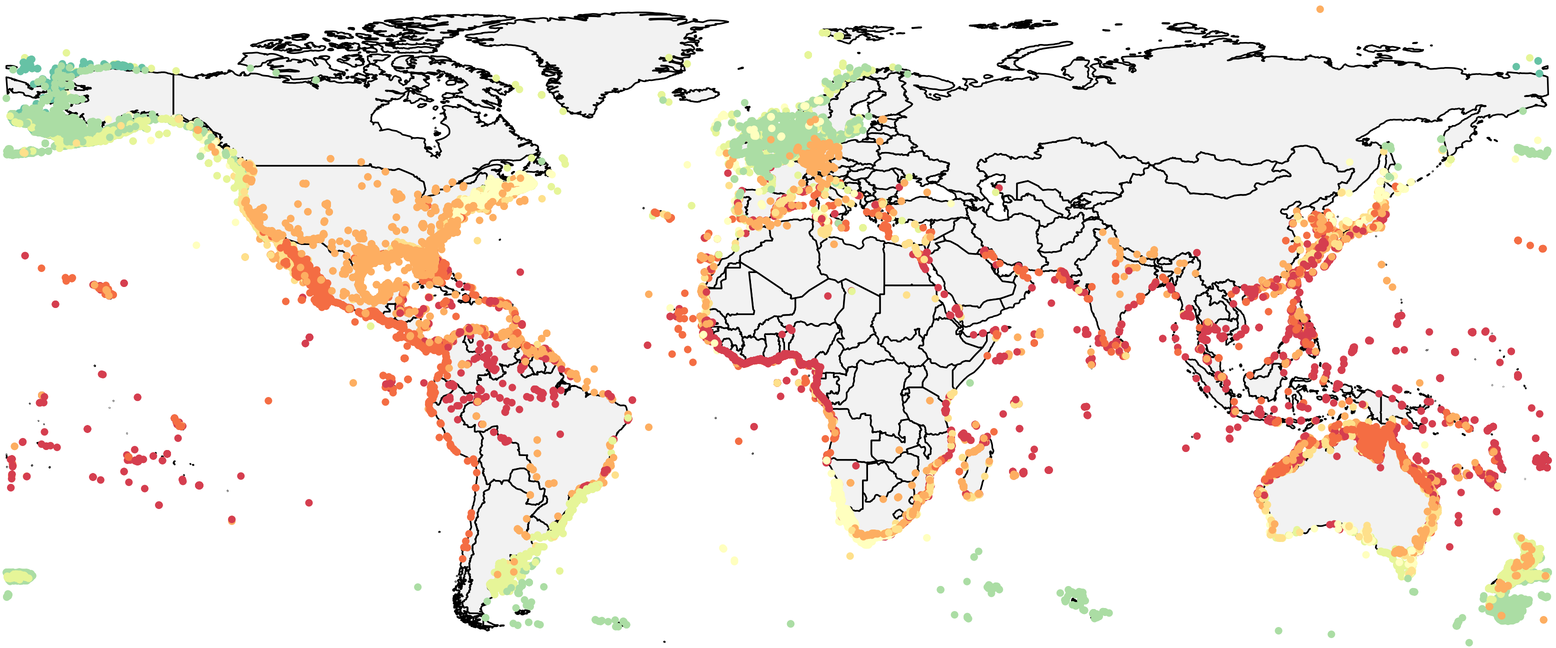

**Figure S3.** Geographic distribution of all 310 flatfish species analyzed in this study. Location data were extracted from GBIF for at most 1,000 individuals from each species, and their latitudes were used to color-code each observation with warm colors at low latitudes, and colder colors at higher latitudes. These same colors are used in Fig. S4 for the species distributed in the northern hemisphere

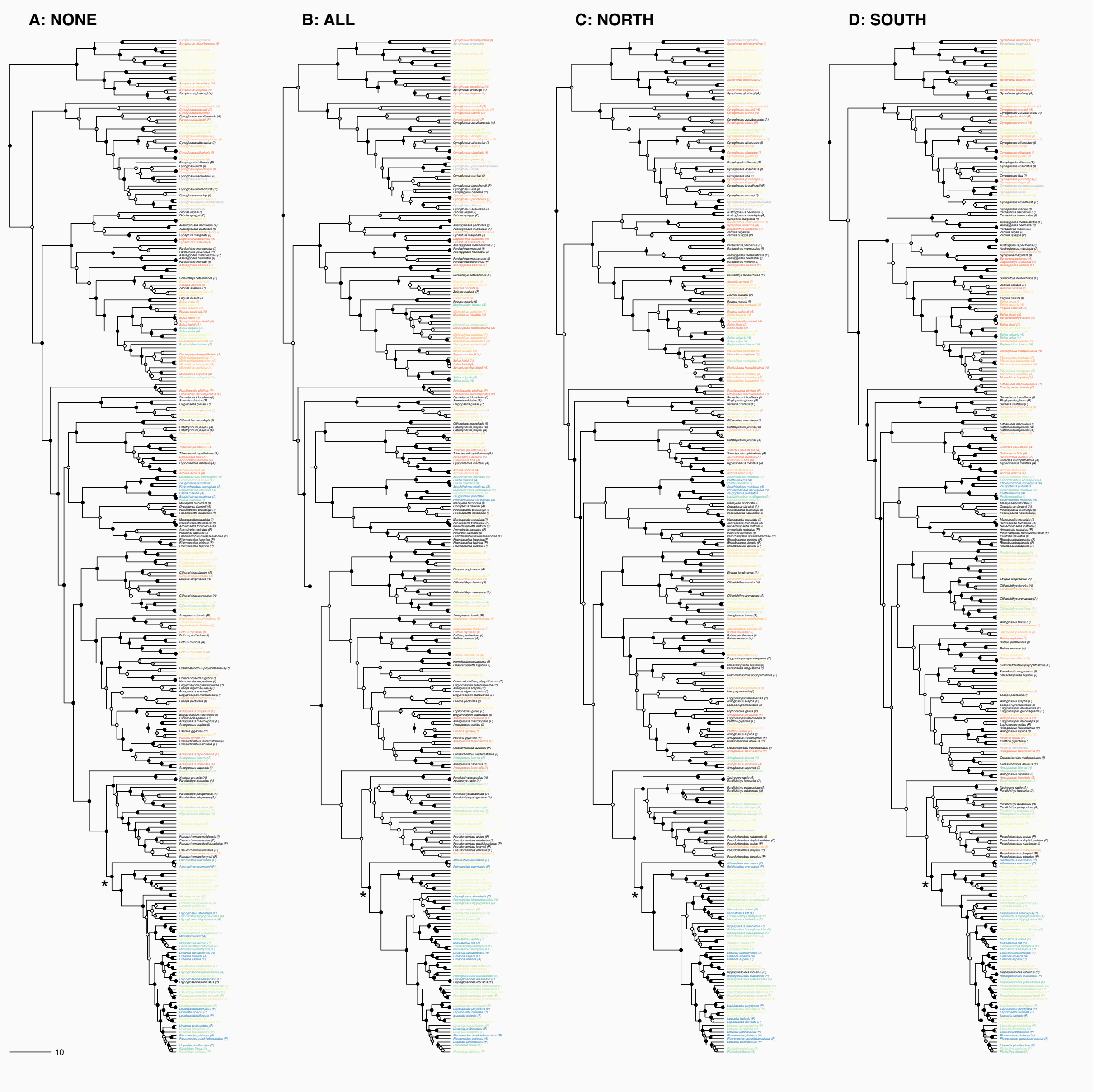


**Figure S4.** Phylogenetic trees excluding Psettodidae, under four different calibration schemes. NONE: no priors were placed on sister taxa; ALL: priors were placed on all pairs of sister taxa; NORTH: priors were placed on only pairs of sister taxa with a northern distribution; SOUTH: priors were placed on only pairs of sister taxa with a southern distribution. Species names are color coded from warm (low latitudes) to cold (high latitudes) for species that are distributed in the Northern hemisphere (those in the Southern hemisphere have their names in black; those with no data in GBIF have their name in light gray). Scale bar: 10 MY. Posterior probabilities are coded as: $Pr \geq 0.9$ (black); $0.7 \leq Pr < 0.9$ (gray); $Pr < 0.7$ (white). The asterisk indicates the northern clade. Approximate locations: Atlantic (A), Indian (I), and Pacific (P) oceans.
